# Prevalence of Staphylococcal Superantigens and their association among bacteremic and Infective Endocarditis patients in Egypt

**DOI:** 10.1101/2020.02.03.932921

**Authors:** Heba M. Elsherif, Zeinab H. Helal, Mona R. El-Ansary, Zeinab A. Fahmy, Wafaa N. Eltayeb, Sahar Radwan, Khaled M. Aboshanab

## Abstract

**Aim:** Infective endocarditis (IE) is a major complication of *Staphylococcus (S.) aureus* infection in humans particularly those with bacteremia. Although *Staphylococcus* species are commensal on or in different parts of the human body, it is also known to be a serious pathogen causing bacteremia and sepsis that could lead to IE. Therefore, our aim was to assess the prevalence as well as phenotypic and genotypic association of the Staphylococcal superantigens (SAgs) among bacteremic and IE patients.

**Methods:** This study was conducted on *Staphylococcus* isolates recovered from bacteremic and IE patients. The isolates were screened phenotypically for the detection of SAgs including Staphylococcal enterotoxins (SEs) and toxic shock syndrome toxin-1 (TSST-1). Molecular detection and analysis of *sea, seb, sec, sed, see* and tsst-1, the major SAgs coding genes were performed using PCR and agarose gel electrophoresis, respectively. The obtained findings were statistically analyzed using standard methods.

**Results:** Detection of SAgs using ELISA revealed that 12 (46%) isolates were positive for enterotoxin production. However, the PCR revealed that 19 (73%) isolates were positive for enterotoxin genes with the highest prevalence of the *sea* gene (79%), followed by the *seb* (63%), tsst-1 (21%). The least frequent gene was the *sed* (5.3%). Accordingly, phenotypic and genotypic screening for prevalence of SAgs among Staphylococcal isolates showed significant difference *(P value* =0.046703), however, no significant correlation could be observed among the coagulase negative Staphylococci (CoNS) isolates (P *value* =0.248213). Statistical correlations between bacteremic and IE isolates with respect to prevalence of SAgs, showed no significant difference (P-value = 0.139, Effect size = 0.572) indicating no specific association between any of the detected SAgs and IE.

**Conclusion:** no significant difference has been found between Staphylococcal IE and bacteremia isolates regarding both phenotypic and genotypic detection of the most commonly SAgs. Therefore, all Staphylococcal bacteremic patients are suspected for IE. Also, future work should be conducted for analysis of SAgs gene expression.

## Introduction

*S. aureus* is a dangerous and versatile human pathogen because of its ability to cause various types of infections, including skin and soft tissue sepsis, pneumonia, bloodstream infections (BSIs), osteomyelitis and infective endocarditis (IE) [1]. Higher mortality rate (15–25%) was recently reported from serious *S. aureus* infections, particularly bacteremia and endocarditis [2,3]. IE is a major devastating complications of Staphylococcal bacteremia [3,4]. In 2018, Asgeirsson *et al.* reported that *S. aureus* is the foremost cause of IE where it comprises about 15–40% of all IE cases worldwide [3].

IE is caused principally with bacteria or fungi where serious clinically relevant complications including, damaging of one or more of the heart valves, necrosis of mural endocardium, or sometimes septal defects could occur [5]. Moreover, IE is distinguished by the development of “cauliflower– like” vegetations, consisted of host factors including, fibrin, platelets and bacterial aggregates on the damaged endothelium of heart valves [6,7]. Host related risk factors for IE have been established. Patient groups at high risk of developing *S. aureus* bacteremia include patients with high rates of colonization, immunosuppressive conditions such as cancer, diabetic patients as well as patients on hemodialysis. Insertion of foreign bodies such as central and peripheral venous catheters, prosthetic heart valves and joints increases the risk of infection [8,9]. However, the involvement of bacterial complications still needs further studies to be identified, as the basis of staphylococcal virulence and switching between commensal and pathogenic phenotypes is still obscure.

Based on the European viewpoint, clinical picture, microbiological analysis and echocardiographic investigations are the most helpful techniques that are usually used for diagnosis of IE [10]. The widely accepted Duke criteria provide high sensitivity and specificity for the diagnosis of IE where series of major and minor clinical and pathologic criteria are implemented [11, 12].

SAgs are significant virulence factors that contributed to variety of pathological conditions, including, pneumonia, soft tissue sepsis, toxic shock syndrome, and IE. They have been found to play a very critical part in the pathogenesis of IE [6,7]. IT was recently reported in 2020, that Sags particularly the *S. aureus* enterotoxins play important role in the induction of asthma of hospitalized patients by inducing IgE production [13]. *S. aureus* strains secrete up to 23 of at least 24 serologically distinct SAgs including TSST-1, SEs and the SE-like (SE-l) [8,14,15]. These SAgs have the distinctive capability to concurrently bind both major histocompatability complex and the T-cell receptor, exerting immune response 20% greater than that of ordinary antigens [16, 17]. This immune response is associated with a substantial discharge of various inflammatory cytokines and several interleukins which could have a direct cytotoxic effect on the endothelial cells [7]. Because of the presences of the adhesion surface molecules located on the *S. aureus,* it becomes able to adhere to the cardiac endothelial cells and direct release the cytokines and therefore, the inflammation of the cardiac muscles are initiated causing IE [18]. Despite of the advancements in the field of therapies and infection control, both the morbidity and mortality rates associated with IE has not declined [7,12]. However, the situation of EI becomes worse up on the emergence of multidrug resistant strains such as methicillin-resistant *S. aureus* (MRSA) [2,19].

According to literature, the prevalence of SAgs and their production among *Staphylococcus* clinical isolates associated with IE, particularly in Egypt, has not been well-defined. For that reason we assessed the prevalence of the Staphylococcal SAgs and their association among bacteremic and IE patients.

## Methods

### Clinical specimens and patient data

Blood specimens were collected from 88 bacteremic patients. The blood specimens were submitted to the Microbiology Laboratory, El-Demerdash hospital Ain Shams University and Ain Shams University Specialized hospital, Cairo, Egypt, for routine culture. Specimens were collected during the period from November 2015 to February 2017. A total of 84 (95.5%) blood specimens showed positive blood culture, of these, 18 (21.4%) specimens were collected from patients diagnosed by the cardiologist to have infective endocarditis based on the Modified Duke criteria [20–22]. Blood specimens were collected from patients having fever and preferably on early admission. Patients without fever or those who have been admitted to the hospital for more than one week have been excluded from our study. The whole study was approved by the Faculty of Pharmacy, Ain Shams University Research Ethics Committee (ENREC-ASU-Nr. 65) where both informed and written consent were obtained from patients or parents of patients after explaining the study purpose.

### Identification and antimicrobial susceptibility testing for the *Staphylococcus* isolates

Identification of Gram positive and Gram negative isolates was performed according to Bergy’s Manual [23]. All Staphylococcal isolates were subjected to susceptibility testing against vancomycin (30 μg); clindamycin (2 μg); gentamicin (10 μg) and ciprofloxacin (5 μg) using modified Kirby Bauer disc diffusion method as recommended by CLSI 2016 guidelines [24]. Phenotypically, MRSA isolates were identified by their resistance to cefoxitin disc (30 μg) as recommended by the CLSI, 2016 [24]. *S. aureus* ATCC^®^ 25923 standard strain was used for the quality control of antimicrobial susceptibility tests

### Phenotypic detection of Staphylococcal enterotoxins using ELISA

The presence of SEs A, B, C, D and E in bacterial supernatants was assessed using VIDAS^®^ Staph enterotoxin II kit (BioMerieux, France), following the manufacturer’s protocol. A result with a test value that is less than the threshold value (< 0.13) indicated that the sample either does not contain SE or the toxin concentration was below the detection limit. On the other side, a result with a test value that is ≥ 0.13 indicated the presence of any type of the enterotoxins.

### Molecular analysis of Staphylococcal SAgs

Genomic DNA purification was carried out using Thermo Scientific GeneJET Genomic DNA Purification Kit (Thermo Scientific, UK), following the manufacturers protocol. As shown in table 1, six pairs of primer were used for the PCR amplification of the SAg genes including, *sea, seb, sec, sed, see* and *tsst* genes, coded for SEs A, B, C1, D, E and TSST-1, respectively.[25]. Each PCR reaction contained 12.5 μl of Dream*Taq* Green PCR Master Mix (2X), 100 pmol/μl of each primer for each gene, 100 nmole of chromosomal DNA and continue up to 25 μl with sterile nuclease-free water. DNA Amplification was performed using a Horizontal Thermocycler (Biometra, Germany), with the following thermal cycling profile: initial denaturation step at 94 °C for 5 min, followed by 35 cycles of denaturation at 95°C for 2 min, annealing at 50°C for 2 min, extension at 72°C for 1 min, followed by final extension at 72°C for 7 min. The PCR products was analyzed using 0.8% agarose gel electrophoresis and verified by DNA sequencing [26]

**Table 1:**
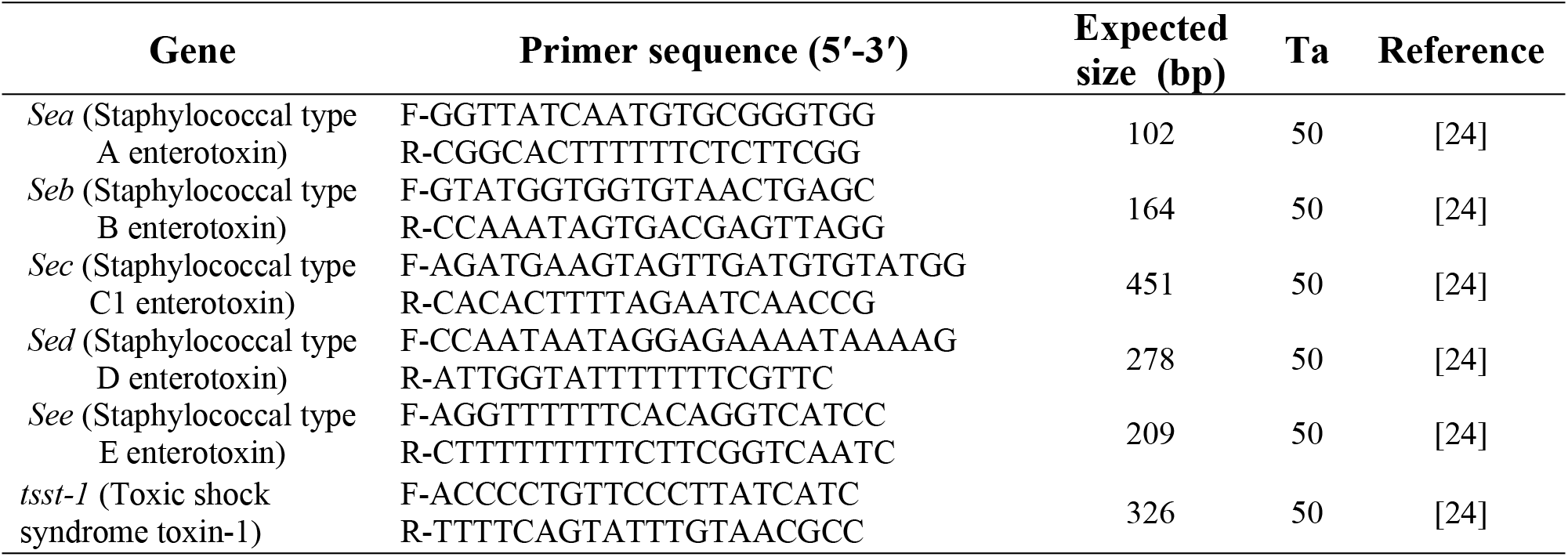
Primer sequences and expected sizes of PCR products

### Statistical analysis

Statistical analysis was performed using Minitab software, version 18.1. Fisher’s Exact test was used for comparisons related to qualitative data. ELISA and total number of detected genes data showed non-parametric distribution, so Mann-Whitney U for the comparisons. A significance level of 0.05 was used.

## Results

### Study population

The study was conducted on 84 positive culture male (65; 77.3%) and 19 (22.6%) female bacteremic patients. Based on the Modified Duke criteria, 18/84 (21.4%) patients were diagnosed by the cardiologist with IE. A summary of these 18 patient’s demographics and clinical characteristics are presented in table 2. The eighteen patients showed native valve endocarditis. There were 14 (77.7%) patients having damage in one single valve, while 4 (22.2%) patients had defects in two valves. Among these patients, the tricuspid valve was the most commonly affected (9; 50%) followed by the mitral (8; 44.4%) and then aortic (5; 27.7%).

**Table 2.** Demographics and clinical characteristics of 18 patients with IE

| Variable | Number | Percentage (%) |
| --- | --- | --- |
| **Gender** |  |  |
| Male | 15 | 83% |
| Female | 3 | 17% |
| **Valve type** |  |  |
| i)-native | 18 | 100% |
| a) Tricuspid (T) | 9 | 50.0% |
| b) Mitral (M) | 8 | 44.5% |
| c) Aortic (A) | 5 | 27.7% |
| ii)-prosthetic | 0 | 0% |
| **Number of Valve affected** |  |  |
| a) Single valve | 14 (77.7%) |  |
| b) Two valves | 4 (22, 2%):<br>T+M (3; 75%)<br>T+A (1; 25%) |  |
| **Age (years)** | 17-48 |  |
| **Smoker** |  |  |
| Yes | 14 | 78% |
| No | 4 | 22% |
| **Addiction** |  |  |
| Injection drug users (IDUs). | 12 | 66.6% |
| **Hepatitis C virus (HCV)** |  |  |
| Yes | 10 | 56% |
| No | 8 | 44% |
| **Surgery performed** |  |  |
| Yes | 4 | 22% |
| No | 14 | 78% |
| **Admitted to ICU** |  |  |
| Yes | 2 | 11% |
| No | 16 | 89% |
| **Comorbidities** |  |  |
| On hemodialysis | 1 | 5.5% |
| Diabetes | - | 0% |
| Cancer | 1 | 5.5% |
| No comorbidities | 16 | 89% |
| **Complications** |  |  |
| Renal failure | 1 | 5.5% |
| Pneumonia/ lung abscess | 2 | 11% |
| Peripheral septic emboli | 1 | 5.5% |
| No complications | 14 | 78% |

### 3.2. Microbial Population and antimicrobial sensitivity

Laboratory examination of 84 positive culture specimens, a total of 85 clinical isolates (83 specimens gave single and 1 specimen was double culture) were recovered, of these, 59 (69.4%), 22 (25.8%) and 4 (4.7%) were Gram positive, Gram negative and *Candida* spp., respectively. Among the Gram positive isolates, the most common organisms identified were the CoNS (30; 50.8%), followed by *S. aureus* (26; 44.1%) and *Streptococcus* spp. (3; 5.1%). The most common CoNS isolated were *S. epidermidis* representing 50%, followed by the *S. lugdunensis, S. haemolyticus* and *S. intermedius* representing 23.3%, 20% and 6.6%, respectively.

As shown in table 3; among the *S. aureus* isolates, 24 (92.3%) were MRSA. Moreover, the sensitivity of *S. aureus* isolates against vancomycin, clindamycin, ciprofloxacin and gentamicin were 92.3%, 65.4%, 61.5% and 50%, respectively. On the other hand, the sensitivity of CoNS against vancomycin, ciprofloxacin and gentamicin clindamycin and cefoxitin, were 93.3%, 40%, 36.6%, 33.3%, and 6.6%, respectively.

**Table 3.**
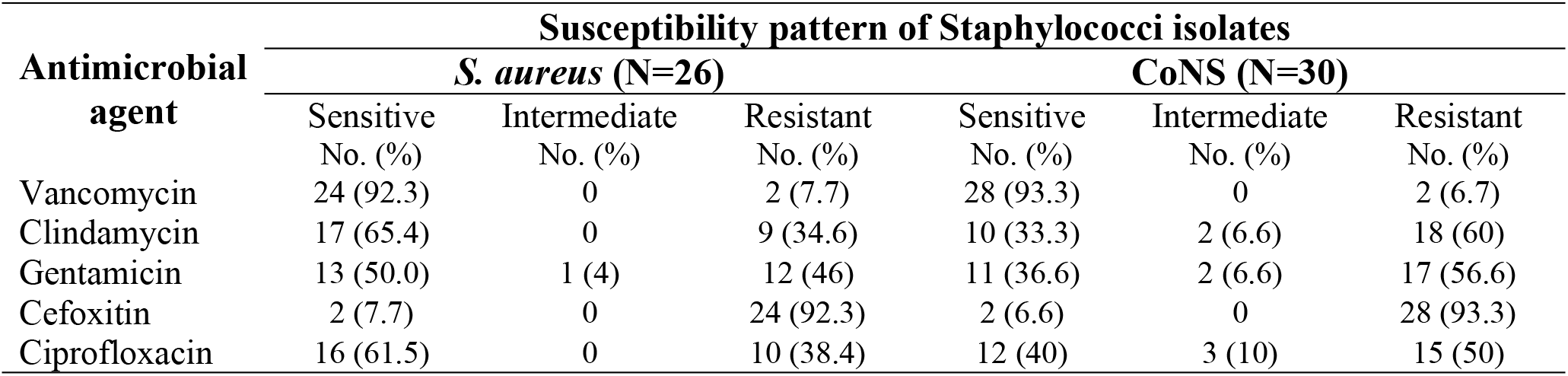
Antimicrobial susceptibility pattern of the recovered *Staphylococci*

### Microbiology of the IE Cases

Out of the 18 blood specimens collected from IE patients, 19 microbial isolates were recovered. The most common pathogen was *S. aureus* (10; 52.6%), followed by CoNS representing (5; 26.3%), Gram negative isolates represented (3; 15.7%) while one isolate was from the *Candida* spp. (5%).

### Phenotypic SAg detection

Twenty six Staphylococcal isolates were selected (15 isolates recovered from IE patients and 11 isolates from bacteremic patients), for the detection of SAgs using ELISA. Only, 12 isolates (46%) were positive SAg production while the remaining 14 isolates (54%) were negative. The 12 positive SAg isolates were, 9 (75%) from IE and 3 (25%) from bacteremic patients without IE. The mean for the ELISA score was 0.995 and the median was 0.05. The P50 was 0.05, which indicated that 50% of the tested isolates showed an ELISA score less than 0.05. The value of P75 was 2.04 which indicated that 75% of the isolates showed an ELISA score less than 2.04, while the P90 was 2.108 which means than 90% of the isolates revealed a score less than 2.108.

### Molecular analysis of Staphylococcal SAgs

As shown in table 4, out of the 26 isolates, 19 (73%) harbored at least one SAg gene, of these, 8 isolates were positive for only one and 11 were positive for two or more genes. Out of the 19 positive isolates, 14 (73.7%) and 5 (26.3%) were from IE and bacteremic patients without IE, respectively. The most frequent gene found among the tested isolates was *sea* gene representing 79% of the isolates, followed by the *seb* and the tsst-1 genes representing 63% and 21%, respectively. The least frequent gene was the *sed* representing only 5.3%. However, *sec* and *see* genes were absent in any of the tested isolates. There was a statistically significant difference in the overall prevalence of the *sea* gene compared to the other genes detected among the tested isolates (*p=0.001*). To contrast differences between *S. aureus* and CoNS genetic profile, among *S. aureus* isolates, *sea* gene was the most prominent gene and was identified in 65% of the studied *S. aureus* isolates, however it was only identified in 33.3% of CoNS. On the other hand, *seb, tsst* and *sed* genes were present in 50%, 20% and 5 % of *S. aureus* isolates, respectively. Though, among CoNS, *seb* and *sed* genes were both present each in 33.3%, while the *sed* gene was not detected in any of the tested isolates.

**Table 4.** Distribution of SAg genes among *Staphylococci*

| Genes | No. (%) of Staphylococcal isolates harbored SAg genes (N= 26) |  |
| --- | --- | --- |
| Detection of gene(s) | Positive: 19 (73%) | EI patients 14 (73.7%);<br>Bacteremic patients without IE, 5 (26.3%) |
|  | Negative: 7 (27%) |  |
| One SAg | 8 (42%) | $P$- value = 0.491 |
| ≥ 2 SAg | 11 (57.9%) |  |
| Types of genes detected |  |  |
| *sea* | 15 (79%) | $P$- value = 0.001 |
| *seb* | 12 (63%) |  |
| *sec* | 0 (0%) |  |
| *sed* | 1 (5.3%) |  |
| *see* | 0 (0%) |  |
| *tsst-1* | 4 (21%) |  |

### Correlation between phenotypic and genotypic detection of SAg among Staphylococci

We studied the statistical correlations between IE and bacteremic isolates with respect to the phenotypic and genotypic detection of SAg. Mann-Whitney Rank Sum Test and Fisher’s Exact test were performed to test for the significance of the SAg as detected phenotypically using ELISA test and genotypically using PCR amplification among bacteremic and IE *Staphylococcus* isolates. As shown in table 5, no significant difference has been found when comparing the results of the ELISA test among IE and bacteremic isolates (P-value = 0.085, Effect size = 0.677). There was no statistically significant difference between prevalence of *sea, seb, sed* and *tsst-1* genes among bacteremic and IE patients (P *value* = 0.426, Effect size = 0.212), (P *value* = 0.453, Effect size = 0.168), (P *value* = 0.423, Effect size = 0.234) and (P-value = 0.113, Effect size = 0.365), respectively. There was also no statistically significant difference has been found between the total number of SAg genes present among IE and bacteremic Staphylococcal isolates (P-value = 0.139, Effect size = 0.572). Statistical analysis indicated significant difference in the genotypic detection method compared to phenotypic detection among the *S. aureus* isolates (P *value* =0.046703). However, there was no significant correlation between genotypic and phenotypic detection among the CoNS tested isolates (P *value* =0.248213).

**Table 5.** Descriptive statistics, results of Fisher’s Exact test and Mann-Whitney U test for comparison between ELISA results and detected genes among bacteremic and IE isolates.

| Outcome | IE (n = 15) | Bacteremia (n = 11) | *P-value* | Effect size |
| --- | --- | --- | --- | --- |
| ELISA [Median (Range)] | 1.92 (0-2.59) | 0 (0-2.05) | 0.085 | *d* = 0.677 |
| Detected genes [n (%)] |  |  |  |  |
| Sea | 10 (66.7%) | 5 (45.5%) | 0.426 | *v* = 0.212 |
| Seb | 8 (53.3%) | 4 (36.4%) | 0.453 | *v* = 0.168 |
| Sed | 0 (0%) | 1 (9.1%) | 0.423 | *v* = 0.234 |
| tsst1 | 4 (26.7%) | 0 (0%) | 0.113 | *v* = 0.365 |
| Total number of genes [Median (Range)] | 2 (0-2) | 1 (0-3) | 0.139 | *d* = 0.572 |

\*: Significant at $P \leq 0.05$

We studied correlation between the ELISA results among staphylococcus isolates in association with the respective genes detected. As shown in table 6, isolates with (*sea*) gene showed statistically significantly higher median ELISA results than isolates without (sea) gene (P-value = 0.005, Effect size = 1.215). There was no statistically significant difference between ELISA results in isolates with and without *(seb* and *tsst1*) genes *(P value* = 0.978, Effect size = 0.010) and (P *value* = 0.940, Effect size = 0.028), respectively. As regards to *sed* gene, no statistical comparison was performed because there was only one isolate harbored this gene.

**Table 6.**
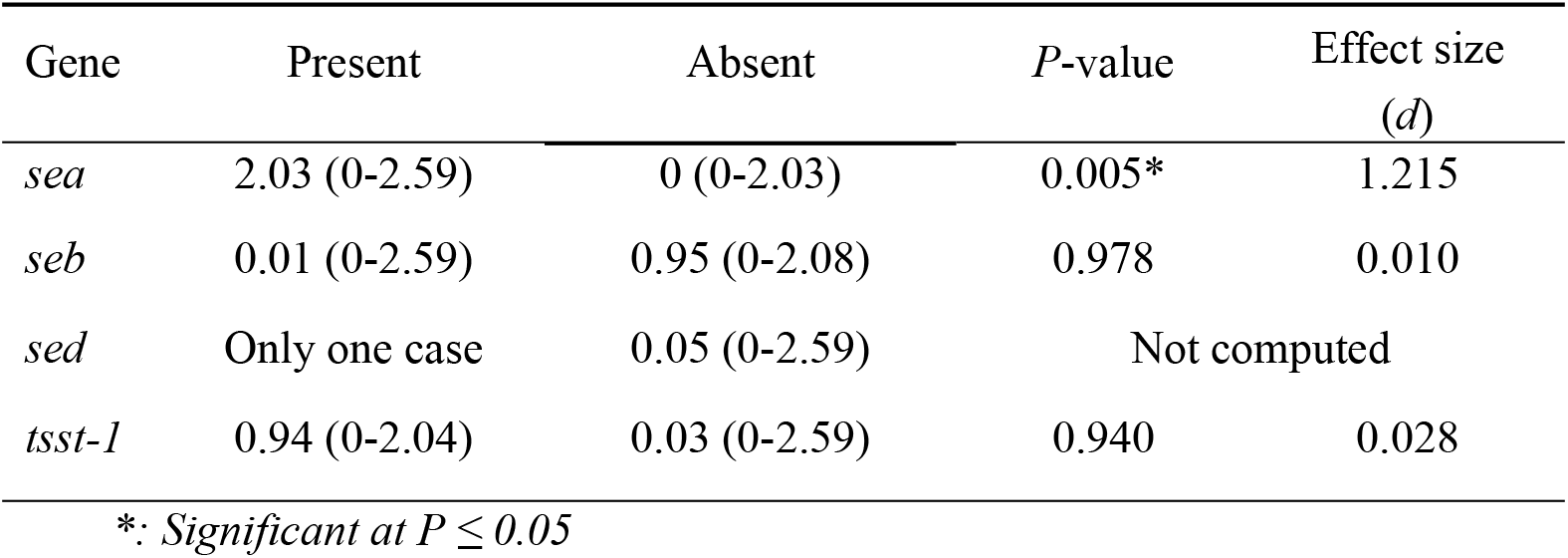
Median, range and results of Mann-Whitney U test for comparison between ELISA results in association with the detected genes.

## Discussion

Globally, BSIs are the major cause of infectious disease morbidity and mortality. Recently, the epidemiology of BSIs has been changed, as a result of many factors, for example increasing globalization, emerging of antimicrobial-resistant organisms, changing population demographic and modifications in health care delivery models [27, 28]. It is reported that *S. aureus* is the second most common species causing BSIs [28]. Genetic variation of genes that encodes for SAg production by *Staphylococcus* spp. may contribute to the occurrence of IE in the course of bacteremia. Therefore, it is important to highlight on the prevalence of SAgs in IE as well as in bacteremic patients.

The cornerstones of clinical diagnosis of IE rely on integration of clinical, microbiological, echocardiography and laboratory findings; these are underlined in the modified Duke criteria for diagnosis of IE. Several studies have reported the high sensitivity of the Duke criteria in the diagnosis of IE [29]. In the present study, the mean age of our patients was 33 ± 11.3 years. The majority of patients were male representing 83% and the remaining 17% were female patients. Similar findings were previously reported [30, 5, 12]. The higher incidence of IE among the young, male patients is attributed to the predominance of the injection drug users (IDUs), which is considered a problem related mainly to the young males in our Egyptian society [31]. The study revealed 75% of the IDUs were HCV positive. These results were in agreement with those reported by a study conducted on IE patients who were intravenous (I.V) addicts; they reported that 86% of the patients were HCV positive [32]. Our findings were in accordance with results reported by a study conducted by Ghosh et al., in which 91% of the patients had fever as the most prevalent symptom [33]. However, absence of fever cannot rule out the diagnosis of IE especially in patients with important clinical features. All our IE patients had native valve IE, this for the reason that postoperative prosthetic valve patients followed a strict follow-up with adequate medical care, so the prevalence of IE among these patients decreased significantly [33]. Among the IDUs, the tricuspid valve was the most commonly affected valve (66.7%). Similar finding was observed in a previous study [34].

The results of our study revealed that the higher prevalence was for the Gram positive isolates (69.4%) causing BSIs compared to the Gram negative isolates (25.8%). Among the recovered Gram positive isolates, the highest prevalence was for the CoNS (50.8%), followed by *S. aureus* (44.1%) and the least prevalence was for the *Streptococcus* spp. (5%). The epidemiology of BSIs towards Gram positive pathogens could be due to the increase in risk factors in the populations including older age, diabetes, end-stage renal disease, intra-cardiac devices, increased use of invasive procedures, I.V drug use as well as frequent insertion of central venous catheters. All these mentioned patient risk factors may lead to the development of complicated BSIs with MRSA as well as CoNS [35, 36].

When assessing the susceptibility of *S. aureus* to different antimicrobial agents, our results revealed that MRSA was responsible for most *S. aureus* bacteremia (92.3%) and also cefoxitin resistance among CoNS was highly noticed with percentage 93.3%. Our results were in agreement with the results of several studies conducted in Egypt reporting the high frequency of MRSA among *S. aureus* isolates with percentages 40% and 88%, respectively [37, 38]. As for the susceptibility of the isolates to vancomycin, 92.3% of the *S. aureus* isolates were sensitive and 93.3% of the CoNS were also sensitive to vancomycin. Accordingly, vancomycin remains the drug of choice, and the most appropriate and commonly used treatment for Staphylococcal BSI’s. In particular, vancomycin is endorsed by the Infectious Diseases Society of America (IDSA) MRSA guidelines as the main treatment choice for MRSA bacteremia [39].

IE usually results from infection by Gram-positive bacteria and infrequently from Gram-negative bacteria. This may be due to that the Gram positive bacteria have capability to adhere and inhabit damaged valves [40]. In addition Gram positive bacteria are armed with numerous superficial adhesins that arbitrate attachment to extracellular host matrix proteins [41]. *S. aureus, Streptococcus* spp., and enterococci are the most common IE pathogens which responsible for more than 80% of IE cases [42, 43]. Historically, *Streptococcus* species have been the main causative microorganisms of IE. However, recently, other pathogens have gained importance. *S. aureus* has become the predominant causative organism in the world, in both hospital settings as well as the community, followed by CoNS [36, 1]. Accordingly, *S. aureus* was the most commonly isolated pathogen (52.6%), followed by CoNS (26.3%). Same finding were reported in a study conducted by Fatima et al., where *S. aureus* was found to be the predominant organism causing IE (38%) [37].

To date, several SAgs have been identified and globally, SAg genes have been found in over 70% of *S. aureus* isolates [44, 45]. Various immunological and molecular methods have been developed for the phenotypic and genotypic detection of SAgs. The prevalence of five SEs encoding genes (A-E) as well as the *tsst-1* was investigated by PCR amplification. The results revealed that among the total Staphylococcal isolates, 57.9% carried two or more genes of the assessed SAgs, while 42% of the isolates had only one gene. From the 15 isolates recovered from IE patients, 93% of the isolates had at least one SAg gene and 53% had two SAg genes. In addition, among *S. aureus* isolates there were 80% of the isolates had at least one SAg gene and these were in accordance with those reported in several studies, where they found that 70-90% of the isolates had one SAg gene [44, 46]. However, lower percentage was reported by Chung et al., where out of the 124 isolates, 63 *S. aureus* isolates (50.8%) had at least one SAg gene [47]. Among the *S. aureus* isolates SAg genes, *sea* was the most commonly found gene followed by the *seb* gene, tsst-1 and the least prevalence was for the *sed.* The genes coding for enterotoxins C1 and E were not found among the tested isolates. Our results were in accordance with other reports [48, 49]. They found the highest frequency for *sea* gene, followed by the *seb* gene and *sed* genes. In contrast to our study, Nhan *et al.,* found that *sec* and *seb* genes were the most prevalent toxin genes in their study [50].

The molecular detection of SAg was found to be more sensitive and efficient than the ELISA test, since the results of the PCR amplification revealed that 20 out of the 26 isolates were positive for SAg genes. However, only 12 isolates were positive for enterotoxin production. This could be due to the low level production of enterotoxins and TSST-1 by some isolates, which are not detected by VIDAS ELISA. Also, it is unable to differentiate between active and inactive SEA [51]. Another possible explanation is that expression of SAgs genes is more prominent *in vivo* than in *in vitro* culture methods.

In spite of this, the question remains as to whether IE *Staphylococcus* isolates differ from non-IE bacteremia isolates. Our results showed no significant difference between Staphylococcal IE and bacteremia isolates with respect to both phenotypic and genotypic detection of the most commonly found SAgs. Our data rule out the possibility of a single specific SAg responsible for the occurrence of IE in the course of Staphylococcal bacteremia. Our results were in accordance with Bouchiat et al., and Gallardo-García et al., where they found no association between any SAg and IE [48, 52]. On the other hand, in 2014, Chung et al. analyzed a series of 124 *S. aureus* isolates in IE, and found a significant correlation between SAgs and IE [47]. Moreover, in 2012, Tristan et al. found that the genes encoding toxic shock syndrome toxin-1 and staphylococcal enterotoxin A, the two major SAgs from *S. aureus,* were enormously widespread in IE isolates from the USA 93.9% and 64.9%, respectively [46]. Accordingly, they suggested that IE isolates carry specific virulence factors that differ from those found in isolates tested from patients suffering other infections [46].

## Conclusion

In conclusion, there was a statistically significant difference between phenotypic and the genotypic detection methods among the Staphylococcal tested isolates. On the other hand, our study revealed that no significant difference has been found between Staphylococcal IE and bacteremia isolates regarding both phenotypic and genotypic detection of the most commonly SAgs. Accordingly, all Staphylococcal bacteremic patients are suspected to have IE. It is important to note that one limitation of the study was unable to establish SAg gene expression *in vitro.* However, detection of SAgs gene expression will be made in our future research.

## ACKNOWLEDGMENT

The research team expresses sincere gratitude to Dr Zeinab Abdelsalam Fahmy, Professor of Cardiology, Faculty of Medicine, Ain Shams University for providing us with the diagnosis of the IE cases. We express also our gratitude to the Microbiology Laboratory, El-Demerdash hospital Ain Shams University and Ain Shams University Specialized hospital, Cairo, Egypt for providing us with the clinical isolates recovered from blood specimens. Special thanks are to Dr. Al Baraa El Said for analyzing the data statistically.

